# Chloroquine Up-regulates Expression of SARS-CoV-2 receptor Angiotensin Converting Enzyme-2 in Endothelial Cells

**DOI:** 10.1101/2024.05.10.593585

**Authors:** Hien C. Nguyen, Shuhan Bu, Lynn Wang, Aman Singh, Krishna K. Singh

**Affiliations:** Department of Medical Biophysics, University of Western Ontario, London, ON, N6A 5C1, Canada; Department of Anatomy and Cell Biology, Schulich School of Medicine and Dentistry, University of Western Ontario, London, ON, N6A 5C1, Canada

**Keywords:** Chloroquine, Autophagy, Endothelial Cells, EndMT, COVID-19, ACE2

## Abstract

**Background and Purpose:** The novel severe acute respiratory syndrome coronavirus-2 (SARS-CoV-2) posed a serious threat to global public health. Hydroxychloroquine (HCQ), which is a derivative of Chloroquine (CQ), was a WHO-recommended drug to treat COVID-19 with mixed effects. The purpose of the present study is to evaluate the plausible mechanisms of HCQ actions behind its observed mixed effect.

**Key Results:** We demonstrate that CQ-treatment significantly up-regulates mesenchymal markers and SARS-CoV-2 receptor ACE2 in cultured endothelial cells.

**Conclusions & Implications:** The detrimental effect of HCQ in seriously ill COVID-19 patients might be due to CQ-induced increased expression of endothelial ACE2 exacerbating the severity of SARS-CoV-2 infection. Our study warrants further investigation in animal models and humans and caution while prescribing HCQ to patients with an impaired renin-angiotensin-aldosterone system, such as in hypertension, cardiovascular diseases, or chronic kidney disease; particularly with ACE-inhibitors or statin therapy.

## Introduction

The coronavirus disease 2019 (COVID-19) pandemic – caused by the novel severe acute respiratory syndrome coronavirus-2 (SARS-CoV-2) – has posed a serious social and economic threat worldwide, infecting over 700 million and killing more than 7 million people. The real course of the disease appears complex and still not fully elucidated. However, a cytokine-surge causing inflammation leading to pneumonia and lung failure, followed by cardiovascular complications with wide-spread coagulation in the lungs and endothelial dysfunction, rapidly emerged as a key threat in COVID-19 [1-3]. Currently, there are no approved drugs with proven clinical efficacy to effectively and completely treat or prevent COVID-19. Previously, the US Food and Drug Administration (FDA) approved the limited emergency use of hydroxychloroquine (HCQ) [4, 5]. While the drug’s positive effects were shown, detrimental effects or no-effect results in seriously-ill patients were also reported [6]. Other drugs such as arbidol, remdesivir, and favipiravir are currently in use to treat COVID-19, with mixed reports on their efficacy in the treatment of COVID-19 [5]. There is a need for an effective drug to completely prevent and treat COVID-19 as well as reduce the SARS-CoV-2 transmission rate (R0) with minimal side-effects.

There is unequivocal evidence that angiotensin-converting enzyme 2 (ACE2) is the ‘entry door’ for SARS-CoV-2, allowing it to infect cells [7, 8]. ACE2, is a key enzymatic component of the counter-regulatory pathway of the renin-angiotensin-aldosterone system (RAAS), which regulates blood pressure, inflammation, and fibrosis [9]. Impaired RAAS is the main mechanism behind the pathophysiology of hypertension, cardiovascular disease, and chronic kidney disease and these conditions are routinely treated with RAAS blockers and statins [9-11]. Evidences indicate that RAAS blockade by ACE inhibitors and statins enhance ACE2 expression in animal models, which, in part, may also contribute towards improved cardiovascular function in patients; however, increased ACE2 expression is expected to be detrimental for COVID-19 patients [10]. In this scenario, evaluation of HCQ on ACE2 expression is of immense importance for all COVID-19 patients, particularly for those with pre-existing pathologies which compel the use of RAAS blockers or statins therapy. ACE2 is mainly expressed on endothelial cells (**EC**), as well as epithelial cells [2, 7]. ECs contribute to more than 30% of all cells in the lungs [12], constitute the innermost layer of every blood vessel, and respond to constantly varying hemodynamics to maintain homeostasis [13]. ECs are plastic and as such they have the capability to lose endothelial characteristics and transition into ‘stem cell-like’ mesenchymal cells, a process known as endothelial-to-mesenchymal transition (EndMT) [14, 15]. EndMT play roles in development and also in disease such as inflammation and fibrosis [16, 17].

Hydroxychloroquine [**HCQ;** a less toxic derivative of Chloroquine (**CQ**)], which is an FDA-approved drug to treat malaria, is an autophagy inhibitor and was tested in various clinical trials against COVID-19 [18]. Autophagy is a key homeostatic process where cytosolic components are degraded and recycled through lysosomes for reuse [18]. Viruses act by hijacking the autophagy pathway for their propagation [19]. We have previously demonstrated that inhibition of endothelial autophagy *via* genetic deletion of autophagy-related gene 7 (ATG7) induces EndMT-like phenotypic switching in ECs [20]. Endothelial-specific loss of autophagy-induced EndMT also exacerbated lung fibrosis in the mouse model [20]. We then very elegantly demonstrated that endothelial-specific loss of autophagy attenuates thrombosis by reducing tissue factor expression in animals *in vivo* [21]. ACE2 is expressed basally and widely on the endothelial cells (ECs), but the effects of CQ on ACE2 expression and EndMT are unknown [2, 7]. Accordingly, we sought to investigate the effects of CQ on EndMT-like phenotypic switching in ECs and the expression of ACE2. For the first time, and also to our surprise, we provide evidence that CQ does not induce EndMT-like phenotypic switching but up-regulates ACE2 expression, which might be the mechanisms behind the mixed effect of HCQ in seriously-ill COVID-19 patients, thus encouraging personalized therapy in COVID-19 patients, particularly with pre-existing diseases due to impaired RAAS signaling and on RAAS-inhibitors or statin therapy.

## Methods

Human umbilical vein ECs (HUVECs, passage 4-6, pooled, Lonza) were cultured in EC growth medium-2 (EGM™-2 Bulletkit™; Lonza) containing growth factors or MCDB 131 (Gibco) supplemented with serum and antibiotics. Following 60-70% confluence, the cells were starved over-night in 1% FBS and then treated with 5, 10, 20, 50 and 100 μM of CQ diphosphate (Sigma). The control groups were treated with diluent. Total RNAs and proteins were extracted 24-hours post-treatment. siRNA-mediated ATG7-silencing was performed in HUVECs with 5mM of siATG7 or scrambled control (Ambion) and the Dharmafect-4 transfection reagent (Dharmacon) in accordance with the manufacturer’s guidelines[22]. Total RNAs and proteins were extracted 48-hours post-transfection [20]. Total RNAs were extracted using Trizol (Invitrogen), and cDNA was synthesized (Quantitect, Qiagen). Quantitative (q)PCR was performed to measure the expression level of genes using SYBR® Select Master Mix (Applied Biosystems) and primers for ACE2 [23], CD31, VE-Cadherin, N-Cadherin, fibroblast-specific protein-1 (FSP-1) and αSMA (alpha smooth muscle actin) in QuantStudio™ 3 Real-Time PCR System [24, 25]. Primer sequences for A Disintegrin And Metalloproteinase Domain 17 (ADAM17) were 5’-GTGGATGGTAAAAACGAAAGCG-3’-forward & 5’-GGCTAGAACCCTAGAGTCAGG-3’-reverse and for Transmembrane Serine Protease 2 (TMPRSS2) were 5’-GTCCCCACTGTCTACGAGGT-3’-forward and 5’-CAGACGACGGGGTTGGAAG-3’-reverse. GAPDH was used as internal control [24]. Fold gene expression was calculated using the 2^−ΔΔCt^ method. Total cell lysates were prepared in RIPA buffer (Sigma), and proteins were extracted and quantified. Equal amounts of protein were loaded on sodium dodecyl sulfate (SDS) polyacrylamide gels for immunoblotting analysis. The primary antibodies were utilized at a dilution of 1:1000 for ACE2 (abcam # ab15348), P62 (Cell Signaling # 5114) and GAPDH (Cell Signaling # 5174). The blots were developed using an enhanced chemiluminescence substrate (SuperSignalTM, Life Technologies) and a ChemiDocTM imaging system (Bio-Rad). The differences between the two groups were calculated using Student’s T-test, and differences between more than two groups were calculated using one-way ANOVA with Tukey’s test. A p-value of <0.05 was considered significant. The data are presented as mean±SD.

## Results

Endothelial function measured with different dose of CQ reached its maximum value at 20 μM [26], which is ∼2-10-fold higher than that measured in plasma with the regimens used to treat rheumatoid disease or Malaria [26-28]. However, due to its weak base properties of CQ, it accumulates >1000-fold in the intracellular acidic compartments within a few minutes, which leads to a very high gradient of CQ within and around the cell [26, 28]. Accordingly, to understand the effect of CQ on EndMT, we treated HUVECs with a range (5, 10, 20, 50 and 100 μM) of CQ doses for 24 hours. We used HUVECs, as it represent the standard cellular model to study endothelial cells *in vitro* [23-25]. Loss of endothelial autophagy is known to induce EndMT [20], therefore, first, to evaluate any effect of CQ on the process of EndMT, ECs were treated with different doses of CQ, and microscopic snapshots were taken after 24 hours of CQ treatment. Notably, the conspicuous EndMT-related changes, where the endothelial typical “cobble-stone” phenotype changes to “elongated, smooth and spindle-shaped” morphology, was not observed for all the doses of CQ treatment to ECs, however, the change was clearly evident in the ATG7-silenced positive-control ECs (**Fig. 1A**). Next, to evaluate EndMT at molecular level, we measured the expression level of endothelial markers; CD31 and VE-Cadherin, and mesenchymal markers; N-cadherin, FSP-1 and αSMA, which did not indicate EndMT but we did observe a significant up-regulation of endothelial markers CD31 and VE-Cadherin and mesenchymal markers N-Cadherin, FSP-1, and αSMA at higher doses (20, 50 and 100μM) of CQ (**Fig. 1B**). Lower doses (5 and 10 μM) of CQ appeared to ineffective (**Fig. 1B**). We then treated ECs with 20, 50 and 100μM of CQ for 24 hours and measured the expression level of ACE2. Our qPCR data demonstrated a significant increase in the expression level of ACE2 in CQ-treated ECs compared to vehicle-treated control ECs (**Fig. 1C**). Our transcript data was further confirmed by immunoblotting, which also showed an increase in the expression level of ACE2 in all CQ-treated ECs in comparison to vehicle-treated ECs (**Fig. 1D**,**E**). CQ is a well-established autophagy inhibitor [21]. Accordingly, we measured autophagy inhibition by measuring the P62 levels, which gets degraded in the process of autophagy but is accumulated when autophagy is impaired [20]. Our immunoblot data demonstrated an increased accumulation of P62, thereby autophagy inhibition, in 20, 50 and 100μM of CQ-treated ECs (**Fig. 1F**,**G**). To understand the role of autophagy *per se* on endothelial ACE2 expression, we genetically inhibited autophagy by silencing ATG7 in ECs and measured ACE2 expression. We successfully silenced ATG7 (**Fig. 1H**) and then performed qPCR for ACE2 in ATG-7 silenced and scrambled control treated ECs. Contrasting to CQ treatment, we observed a significant down-regulation of ACE2 in ATG7-silenced ECs in comparison to control ECs (**Fig. 1I**), indicating, if any, the existence of an indirect and autophagy independent role of autophagy in CQ-induced expression of ACE2 in ECs. Furthermore, TMPRSS2 and ADAM17 both cleave ACE2, which is a critical step for SARS-CoV-2 cellular entry [29, 30]. Accordingly, next we measured the expression levels of TMPRSS2 and ADAM17 following 20, 50 and 100μM CQ treatment to ECs and observed a significant down-regulation, in comparison to vehicle treated-ECs (**Fig. 1J, K**).

**Figure 1.**
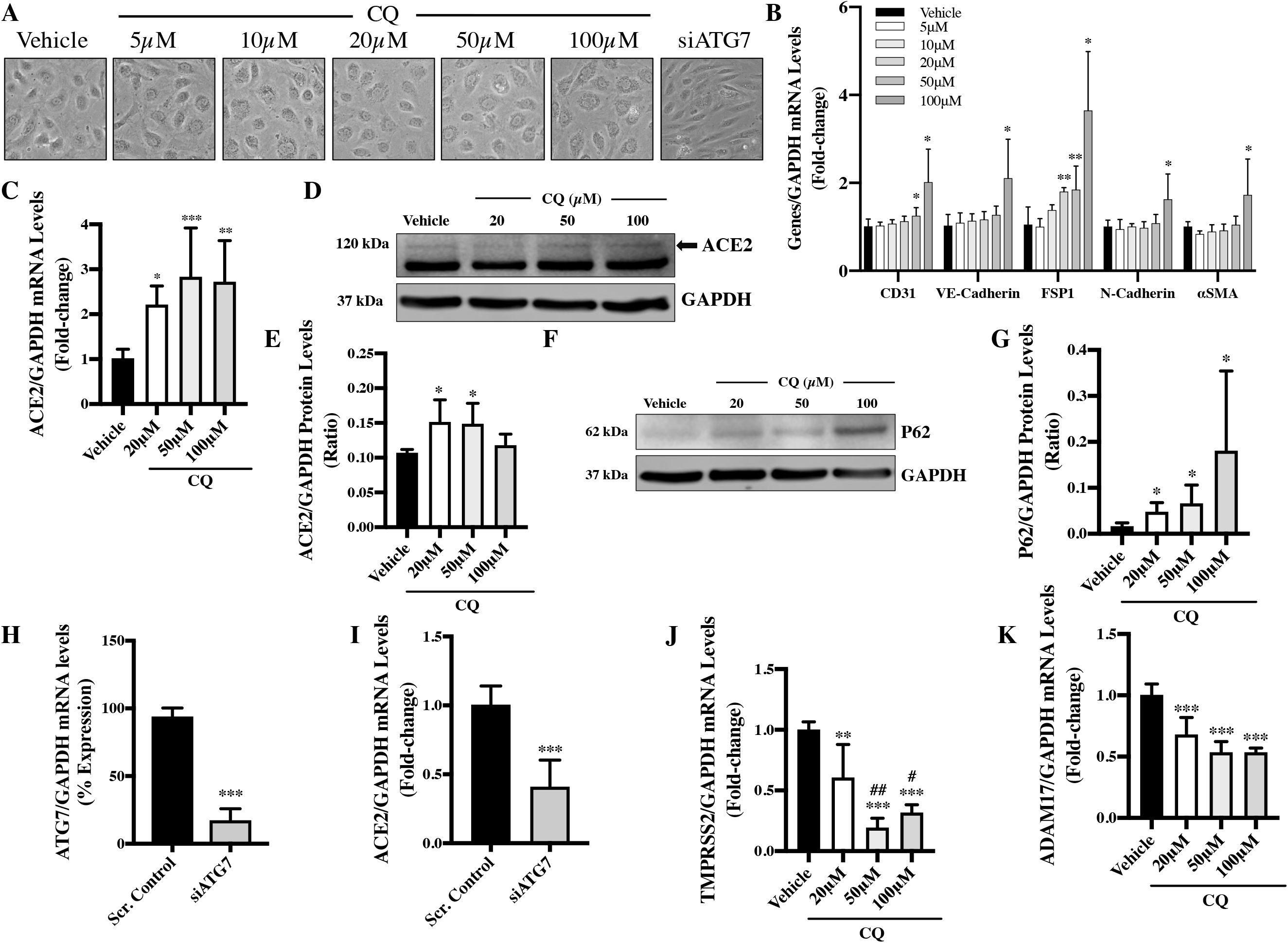
CQ-treatment does not induce EndMT but up-regulates ACE2 expression in ECs. HUVECs were treated with either vehicle-control or 5, 10, 20, 50 and 100μM of CQ for 24 hours and then visualized under light microscope (20X) for **(A)** morphological changes. ATG7-silenced ECs were used as a positive control for EndMT. Later RNA and protein were extracted from CQ-treated HUVECs to perform **(B)** qPCR for EndMT markers, **(C)** ACE2 and **(D)** immunoblot and **(E)** quantification for ACE2, **(F)** immunoblot and quantification for P62 **(G)**. GAPDH was used as a loading control for immunoblotting. HUVECs were transfected either with scrambled control or siATG7 for 48 hours and total RNA were extracted to perform qPCR for **(H)** ATG7, and **(I)** ACE2. HUVECs were treated with either vehicle-control or 5, 10, 20, 50 and 100μM of CQ for 24 hours and then RNA were extracted to perform qPCR for **(J)** TMPRSS2 and **(K)** ADAM17. *p < 0.05; **p < 0.01; ***p < 0.0001 *versus* corresponding scrambled or vehicle control group. #p < 0.05; ##p < 0.01 *versus* corresponding 20μM CQ group. N = 4-6 independent experiments in triplicates for RNA and duplicate or triplicates for proteins. The data are presented as mean±SD.

## Discussion

Chloroquine has been safely used to treat malaria for over 200 years. It was also an approved treatment option against the COVID-19 pandemic, albeit with mixed clinical outcomes. There are reports suggesting beneficial as well as detrimental effects of CQ in seriously-ill COVID-19 patients. Based on the data from the late clinical trials, including the Solidarity Trial, the UK’s Recovery trial, and the French Discovery trial, the World Health Organization (WHO) announced that HCQ does not result in the reduction of mortality of hospitalised COVID-19 patients. However, CQ is known to have anti-viral effects by increasing the late endosomal and lysosomal pH, resulting in impaired release of the viral genetic material into the cell from the endosome or lysosome, hindering viral replication [31-33]. By this mechanism, CQ may provide a preventive strategy against viral propagation in the case of SARS-CoV-2 infection. HCQ have been employed with significant effects in COVID-19 patients, but detrimental effects of the drug, mainly in seriously-ill patients, were reported. For this reason, HCQ applications were officially halted by WHO. Further investigations into HCQ’s mechanisms of action against COVID-19 are therefore warranted, especially to assist clinical prescription in weighing the beneficial-versus-detrimental side-effects in different COVID-19 patients. Cardiovascular complications, including wide-spread endothelial dysfunction, micro-thrombus formation, and endotheliitis, emerged as the key threats in COVID-19 patients in addition to respiratory disease [1, 2]. The endothelium plays critical roles in a multitude of cardiovascular complications, as well as inflammation and thrombosis [34]. ECs are plastic in nature and carry the potential to lose endothelial characteristics and transition into ‘stem cell-like’ mesenchymal cells, a process known as EndMT [14, 15]. Autophagy inhibition by genetic deletion of autophagy essential gene ATG7 in ECs is known to induce EndMT [20]. EndMT is known to be physiologically involved in development [31] and pathologically involved in several diseases such as pulmonary vein stenosis and anomalous vascular remodeling, as well as cerebral cavernous malformations, cancer progression, and organ fibrosis [16, 35]. Most importantly, it was reported that 16% of lung fibroblasts originated from ECs through EndMT in the pulmonary fibrosis mouse model [36], and loss of endothelial autophagy induced EndMT that exacerbated lung fibrosis in the same model [20]. On this notion, we treated ECs with different doses of CQ and evaluated EndMT microscopically as well as through characteristic molecular markers to understand the effect of CQ on EndMT. To our surprise, we did not observe EndMT-like phenotype switching in CQ-treated ECs (**Fig. 1A, B**). However, we did observe an up-regulation in the expression level of both endothelial and mesenchymal markers upon higher doses of CQ, so further investigation into the effect of CQ in relation to SARS-CoV-2 infection and endothelial function needs to be performed.

SARS-CoV-2 uses the ACE2 receptors for cell entry, which are mainly expressed on ECs [2, 7, 8]. Accordingly, we first evaluated the effects of CQ on ACE2 expression in ECs. Our data demonstrate significant induction of ACE2 expression in ECs by CQ treatments (**Fig. 1C-E**), which is particularly of clinical interest for the COVID-19 patients and their CQ-treatment outcomes. SARS-CoV-2 infection is reported to be enhanced and diminished by ACE2 over-expression [37] and inhibition [26-28], respectively. Our CQ data, hence, showed that HCQ can exacerbate the rate of SARS-CoV-2 infection and severity of diseases by increasing the expression levels of ACE2, a detrimental effect to consider apart from the beneficial autophagy inhibition by CQ in infected patients. ACE2 is a key counter-regulator of (RAAS), and RAAS blockade by ACE inhibitors or statins are the main treatment option for patients with hypertension, cardiovascular disease, and chronic kidney diseases [9-11]. RAAS blockade by ACE inhibitors and statins are known to enhance ACE2 expression, which, in part, contributes towards the cardiovascular benefit in these patients; however, increased ACE2 expression – either due to HCQ or ACE-inhibitors – is expected to be detrimental for COVID-19 patients [10]. It is interesting to note that genetic inhibition of autophagy by deleting ATG7 unexpectedly reduced ACE2 expression (**Fig. 1H**,**I**), which goes along with the process of loss of ATG7-asscociated EndMT, which is associated with reduced expression of most endothelial signature genes [20].

Pharmacologic inhibition of autophagy by CQ and genetic inhibition of autophagy by ATG7 deletion both are known to inhibit LDL degradation and cause LDL accumulation in the endothelium, in turn exacerbating leakage of oxidized LDL into the sub-endothelial spaces inducing oxidative stress, apoptosis, inflammation, and endothelial dysfunction. All of these factors contribute towards enhanced atherosclerosis *in vivo* [38]. Vascular smooth muscle cell (SMC)-specific loss of autophagy in mice *in vivo* exaggerated Angiotensin II-associated adverse aortic remodeling and appreciable cardiac failure-associated mortality [39]. Interestingly, while exacerbated lung fibrosis, atherosclerosis, and adverse aortic remodeling were shown in mice with endothelium-specific and SMC-specific autophagy deficiency, these mice were completely normal and healthy at baseline with no stress [20, 38, 39]. Similarly, while HCQ-mediated autophagy inhibition might be beneficial to otherwise healthy COVID-19 patients without pre-existing pathologies, it might be detrimental in COVID-19 patients with pre-existing cardiovascular complications. Furthermore, TMPRSS2 and ADAM17 both cleave ACE2, but it was recently demonstrated that only the cleavage by TMPRSS2, but not by ADAM17, augments SARS-coV-2 cellular entry [29]. The relationship between CQ, endothelial ACE2 and SARS-CoV-2 infection is further complicated by the observed reduced expression of TMPRSS2 and ADAM17 in ECs following CQ treatment (**Fig. 1K, L**).

Our novel *in vitro* findings about the mechanism of action of CQ provide a plausible explanation for the drug’s mixed effects that have been reported in its uses to treat and prevent COVID-19. Our findings have few limitations, such as use of only one type of EC, lack of SARS-CoV-2 data in ECs and also lack of *in vivo* data, warranting further investigations. Overall, our manuscript provides a better source of necessary knowledge on CQ for researchers and health care practitioners to understand the mixed-effect of HCQ in COVID-19 patients and accordingly plan the future research in this area.

## Acknowledgement

This work was supported by a grant from the Heart and Stroke Foundation of Canada (G-17-0018688) to KS. KS is also the recipient of the 2018/19 National New Investigator Award-Salary Support from the Heart and Stroke Foundation of Canada, Canada.

## Author Contribution Statement

KS conceived and designed the study. KS, HN, SB and LW performed the study. KS, HN, SB and AS helped with data analysis, figures, discussion and manuscript writing.

